# Oxidative Stress and Biomarker Responses in the Atlantic Halibut After Long Term Exposure to Elevated CO_2_ and a Range of Temperatures

**DOI:** 10.1101/510818

**Authors:** B Carney Almroth, K Bresolin de Souza, E Jönsson, J Sturve

**Affiliations:** Department of Biological and Environmental Sciences, University of Gothenburg, PO Box 463, SE-405 30 - Gothenburg, Sweden

**Keywords:** oxidative stress, carbon dioxide, ocean acidification, temperature, climate change, teleost fish, Atlantic halibut, *Hippoglossus hippoglossus*

## Abstract

Oceans are warming and pH levels are decreasing as a consequence of increasing levels of dissolved CO_2_ concentrations. The CO_2_ emissions are predicted to be produce in greater and faster changes in the ocean than any other event in geological and historical records over the past 300 million years. Marine organisms will need to respond to multiple stressors but the potential consequences of global change-related effects in fish are not fully understood. Since fish are affected by many biotic and abiotic environmental variables, including temperature and CO_2_ fluctuations, it is critical to investigate how these variables may affect physiological and biochemical processes. We investigated the effects of elevated CO_2_ levels (pH of 8.0, which served as a control, or 7.6, which is predicted for the year 2100) combined with exposure to different temperatures (5, 10, 12, 14, 16, and 18 °C) in the Atlantic halibut (*Hippoglossus hippoglossus*) during a three month experiment. We assessed effects on antioxidant and cholinesterase enzymes (AChE and BChE), and CYP1A enzyme activities (EROD). The treatments resulted in oxidative stress, and damage was evident in the form of protein carbonyls which were consistently higher in the elevated CO_2_-treated fish at all temperatures. Analyses of antioxidant enzymes did not show the same results, suggesting that the exposure to elevated CO_2_ increased ROS formation but not defences. The antioxidant defence system was insufficient, and the resulting oxidative damage could impact physiological function of the halibut on a cellular level.

## Introduction

The release of carbon dioxide (CO_2_) into the atmosphere is changing the ocean’s chemistry at a pace never before seen. The oceans are becoming warmer and pH levels are decreasing as a consequence of increasing levels of dissolved CO_2_ (Solomon et al., 2009; Steffen et al., 2015). The CO_2_ emissions predicted for the coming centuries are expected to produce greater and faster changes in the oceans than any other event recorded in geological and historical records over the past 300 million years (Caldeira and Wickett, 2003). Since many marine animals have evolved to cope with changes within a certain range of temperature and CO_2_ concentrations, climate change is expected to challenge their ability to function optimally at conditions outside of their scope of tolerance (Portner, 2010). In aquatic ectotherms such as fish, environmental temperature is a crucial variable since it has a direct effect on all biological processes, such as metabolism (Portner et al., 2006) and enzyme kinetics (Kavanau, 1950).

The effects of temperature changes and lower pH have been studies in marine animals and we are beginning to understand consequences and mechanisms involved (Gräns et al., 2014a; Gräns et al., 2014b). For example, recent studies have shown indications of oxidative stress in marine animals exposed to changes in temperature: long term exposure to increasing temperatures induces oxidative damage in lipids in rock fish (*Nothonei sp*.) (Klein et al., 2017); short-term elevation in temperature increases oxidative stress in Antarctic vertebrates and invertebrates (Abele and Puntarulo, 2004); low temperatures are reported to increase oxidative stress in gilthead sea bream (*Sparus aurata*) liver (Ibarz et al., 2010). The Antarctic fish species bald notothen (*Pagothenia borchgrevinki*) responded to acutely increased temperature with an increase in antioxidant defences while long-term temperature increase resulted in oxidative damage (Carney Almroth et al., 2015). Elevated CO_2_ concentrations are also known to increase oxidative stress in cold water species: the great spider crab (*Hyas araneus*) showed up-regulation of genes associated with the detoxification of H_2_O_2_ (ascorbate peroxidase, glutathione peroxidase) (Harms et al., 2014); Pacific oyster larvae (*Crassostrea gigas*) expressed higher levels of five of the six investigated antioxidant proteins (Tomanek et al., 2011a); and levels of the antioxidant protein glutaredoxin were up-regulated in the Sydney rock oyster (*Saccostrea glomerata*) exposed to elevated CO_2_ levels (Thompson et al., 2015).

Fewer studies have exposed fish to multiple stressors. Gräns et al. (Gräns et al., 2014a) show that increased temperature and CO2 affected growth in halibut, but did not determine mechanisms of these effects. Pementel et al (Pimentel et al., 2015) showed that the combined stress of OA and warmer temperatures resulted in the accumulation of peroxidative damage in a flatfish, and that early developmental stages are more susceptible to oxidative stress. The physiology of croaker fish (*Argyrosomus regius*) was found to be impacted by these factors as well, with some changes in oxidative stress parameters (Sampaio et al., 2018). These fish also demonstrated some ability to maintain physiological homeostasis in the face of a third stressor (mercury exposure), but the biochemical repercussions of the physiological responses were not fully understood and warrant further investigation.

Oxidative stress is commonly addressed in these studies since it is an essential physiological mechanism known to be affected by biotic and abiotic factors; it is a process initiated by the imbalance between the production of oxidants and their removal by antioxidants and antioxidant enzymes. Reactive oxidants including reactive oxygen species (ROS), are produced during normal cellular respiration in mitochondria, or leaked from enzymatic activity including the Phase I detoxification enzyme CYP1A, and are normally metabolized by antioxidants. This includes such enzymes as superoxide dismutase (SOD), catalase (CAT), and glutathione peroxides or molecular antioxidants like glutathione (Ozcan and Ogun, 2015). ROS levels are important in homeostasis, and are a key regulator of biological processes, but ROS can also initiate oxidative cascades causing severe cellular damage to proteins, lipids and DNA (Kohen and Nyska, 2002). Increased ROS production is a common consequence of metabolic and acid-base disturbances in animals (Tomanek, 2011a; Tomanek et al., 2011a) and also a common mechanism of toxicity (Valavanidis, 2006). Changes in the concentration of antioxidants or oxidative damage products are often used as indicators of environmental stress and pollutant exposure (Carney Almroth et al., 2008; Carney Almroth et al., 2005; Kohen and Nyska, 2002).

Hepatic function is associated with performance and maintenance of numerous physiological mechanisms, such as metabolism, degradation of endogenous compounds, and detoxification of various substances. Hence, hepatic tissue contains high levels of antioxidant enzymes and antioxidants and enzymes in the liver can be used to assess liver function. In addition, liver tissue is often target for analyses addressing environmental stressors: the measurement of ethoxyresorufin-*O*-deethylase (EROD) activity in fish liver is a well-established indicator of exposure to aromatic hydrocarbons (Förlin et al., 1994). However, induction of CYP1A is also closely related to detrimental effects such as apoptosis (Whyte et al., 2000) and may be influenced by a large number of biotic and abiotic factors, which in turn affects biotransformation processes (Rahman and Thomas, 2012; Whyte et al., 2000).

Changes in abiotic parameters in the environment are also know to affect the catalytic activity of acetylcholinesterase (AChE) and butyrylcholinesterase (BChE) (Pfeifer et al., 2005; Pretti and Cognetti-Varriale, 2001), enzymes important hydrolysis of choline esters in central and peripheral nervous systems or in plasma and tissues. Inhibition of the catalytical activities of these enzymes often used as biomarkers for neurotoxic compounds including pyrethroids and organophosphorus insecticides in fish (Fulton and Key, 2009; Mushigeri and David, 2005). Both AChE and BChE have been indicated as being involved in immunity via modulation of the cholinergic anti-inflammatory pathway (Pohanka, 2014) and BChE is important in regulation of ghrelin, a peptide hormone involved in regulation of appetite and growth hormone secretion (Brimijoin et al., 2016). Several studies have indicated that AChE and BChE are affected by warming temperatures as well as seawater pH in mussels (Pfeifer et al., 2005; Wu et al., 2016).

As the global climate changes, exposing organisms to environmental variations which are predicted to increase in the future, it is crucial to investigate how they may affect physiological processes (Parmesan and Yohe, 2003). Fish will be affected by a multitude of environmental variables, including both temperature and CO_2_ variations. In order to investigate the effects of elevated CO_2_, at levels predicted for the near future, in combination with different temperatures, we conducted an experiment exposing Atlantic halibut to a range of temperatures and water pH for three months. We analyzed the activities of the antioxidant enzymes superoxide dismutase (SOD), catalase (CAT), glutathione reductase (GR), glutathione S-transferase (GST) and glutathione peroxidase (GPx), the levels of protein oxidation measured as protein carbonyls (PC), activities of acetylcholinesterase (AChE) and butyrylcholinesterase (BChE), and also the activity of the phase I detoxification enzyme CYP1a, measured as EROD activity.

## Materials and Methods

### Chemicals and reagents

7-ethoxyresorufin, glutathione reductase (GR), reduced and oxidized glutathione, 5,59-dithiobis (2-nitrobenzoic acid) (CDNB), 1-chloro-2,4-dinitrobenzene (DTNB), pyruvic acid, glucose-6-phosphate, oxidized and reduced nicotinamide adenine dinucleotide phosphate (NADPH), reduced nicotinamide adenine dinucleotide (NADH), butyrylthiocholine, acetylthiocholine iodine, 2,4-dinitrophenyhydrazine (DNPH), ethyl acetate, guanidine hydrochloride, digitonin and protease inhibitor were obtained from Sigma Aldrich (St. Louis, MO, USA). Hydrogen peroxide is from Fluka (Buchs, Switzerland), ethylenediaminetetraacetic acid (EDTA) from Merck.

### The fish model

The Atlantic Halibut (*Hippoglossus hippoglossus*) is a benthic marine fish widely distributed in the northern regions of the Atlantic Ocean and in parts of the Arctic Ocean (Haug, 1990). Juveniles stay in coastal areas of at depths of 20-60 m before migrating to more distant areas of both shallow and deep waters (Glover et al., 2006). The Atlantic Halibut was chosen as a model for our studies because of its large socio-economic and ecological importance, and its wide distribution in the Northern hemisphere, and the optimal and suboptimal temperatures for this species have been described (Gräns et al., 2014a). Since the Arctic marine biomes are warming twice as fast as the global average (Fossheim et al., 2015), and both coastal waters and deep-sea waters (with low water flow; (Cai et al., 2011)) are acidifying faster than marine open waters, the Atlantic halibut physiology and distribution are susceptible to impact.

### Experimental setup and fish treatments

The experiment was part of a larger study and a detailed description of the set-up, fish exposure, water quality, data monitoring and sampling methods are available in previous publications (Bresolin de Souza et al., 2014a; Gräns et al., 2014a). Juvenile fish from both sexes (weight of 16.4 g +/- 0.2 g (SEM), from Fiskey’s hatchery (Þorlákshöfn, Iceland) were exposed for 96 days to six different temperatures (5, 10, 12, 14, 16, and 18 °C). For each temperature treatment, two tanks were supplied with water at current seawater pH and two tanks with reduced pH. The CO_2_ treatments represent the present day *p*CO_2_ 400 μatm (~pH 8.0) and 1000 μatm (~pH 7.7), according to pH predictions for the end of this century (Gräns et al., 2014b; Solomon et al., 2007). The fish were distributed into twelve fish tanks (100 L), each supplied with aerated flow-through seawater from header tanks (200 L), supplied in turn with flow-through seawater from 32 m depth (32.0 ± 0.14 ppt). Fish were kept under a 12 h:12 h light:dark photoperiod, and were fed once a day with 2.5 % of body mass with commercial fish feed. Water parameters were monitored daily in the header tanks and fish tanks. Temperature and salinity were continuously recorded.

### Sampling

Fish were killed with a sharp blow to the head and liver samples were collected from eight fish from each treatment (total of 96 fish). Livers were divided into sub-fractions, frozen in liquid nitrogen, and stored at -80 °C prior to handling. Practices concerning methods of animal handling, exposure, and sampling were approved by the Animal ethical committee Gothenburg, Sweden, (ethical permits 221-2010 and 329-2010).

### Sample preparation

Liver cytosol fractions were prepared according to (Forlin, 1980). Liver samples were homogenized (glass/Teflon) in 4 volumes of 0.1 M Na/K-PO_4_ buffer containing 0.15 M KCl at pH 7.4. Homogenates were centrifuged in two steps, first at 10 000 g for 20 min at 4 °C, and then the supernatant was re-centrifuged for 105 000 g for 1 h at 4 °C. The supernatant (cytosolic fraction) was aliquoted and stored at -80 °C prior to analysis. The cytosolic fractions were used to determine the activities of the antioxidant enzymes. The pellets containing the microsomes were re-suspended in homogenizing buffer containing 20 % glycerol and stored at -80 °C prior to analysis.

For protein carbonyl analysis liver samples were homogenized in 4 volumes of 50 mM phosphate buffer (pH 7.4) containing 1 mM EDTA, 0.1 % digitonin, and a cocktail of anti-proteases (Sigma P8340). Samples were then centrifuged for 20 min at 10 000 g at 4 °C. Supernatants were collected for use in DNPH reactions, and total protein content was determined according to Lowry (Lowry et al., 1951).

### Biochemical analysis

Catalase (CAT, EC 1.11.1.6) activity was measured according to Cribb et al. (Cribb et al., 1989a) using hydrogen peroxide as substrate. Glutathione S-transferase (GST, EC 2.5.1.18) activity was measured according to Stephensen et al. (2002a) using CDNB as substrate. Superoxide dismutase (SOD, EC 1.15.1.1) activity was measured using a SOD assay kit from Sigma Aldrich, according to the manufacturer’s instructions. Glutathione peroxidase (GPx, EC 1.11.1.9) activity was measured by the method of Greenwald (Greenwald, 1985), modified by Stephensen et al. (Stephensen et al., 2002b). Glutathione reductase (GR) activity was measured according to the method described by Cribb (Cribb et al., 1989b). Protein carbonylation (PC) was measured via reaction with DNPH followed by TCA precipitation as described previously (Levine et al., 1994; Reznick and Packer, 1994). All assays were performed on a microplate reader (Molecular Devices) at room temperature. Acetylcholinesterase (AChE, EC 3.1.1.7) activity was measured according to a modification of the spectrophotometric method described by Ellman et al. (1961) adapted to a microplate reader. Butyrylcholinesterase (BChE, EC 3.1.1.8) activity was measured as described for AChE with butyrylthiocholine as substrate instead of acetylthiocholine iodine. Ethoxyresorufin O-deethylase (EROD) activity was measured in the liver microsomal fraction according to the method described by Förlin et al. (Förlin et al., 1994) using rhodamine as standard. Total protein content in cytosol and microsomes was measured using the BCA kit from Pierce, according to the manufacturer’s protocol, using bovine serum albumin as standard.

### Statistics

Data was compared using two-way ANOVA (α = 0.05) with temperature and pH as fixed factors. Tukey Post-Hoc test (95% confidence) was applied. Statistical analyses, figures and tables were prepared using GraphPad Prism 7.00, and probability for Type-I error was set to 5% for all tests. All data that did not display homogeneity of variance according to Levene’s test, or normal distribution according to the Shapiro-Wilk test, were log-transformed prior to testing. T-tests were used to assess differences between duplicate aquaria; no differences were found so the samples both tanks within a treatment were pooled t for subsequent analyses.

## Results

In the present study the oxidative stress indicators (SOD, CAT, GR, GST, GPx, and PC), esterases AChE and BChE, and phase I detoxification activity (EROD) were measured in liver of Atlantic halibut. Results indicate the occurrence of oxidative stress in the elevated CO_2_-treated fish, though temperature also appears to play an important role in the balance of antioxidant homeostasis. Results are displayed in Figures 1, 2 and 3.

**Figure 1.**
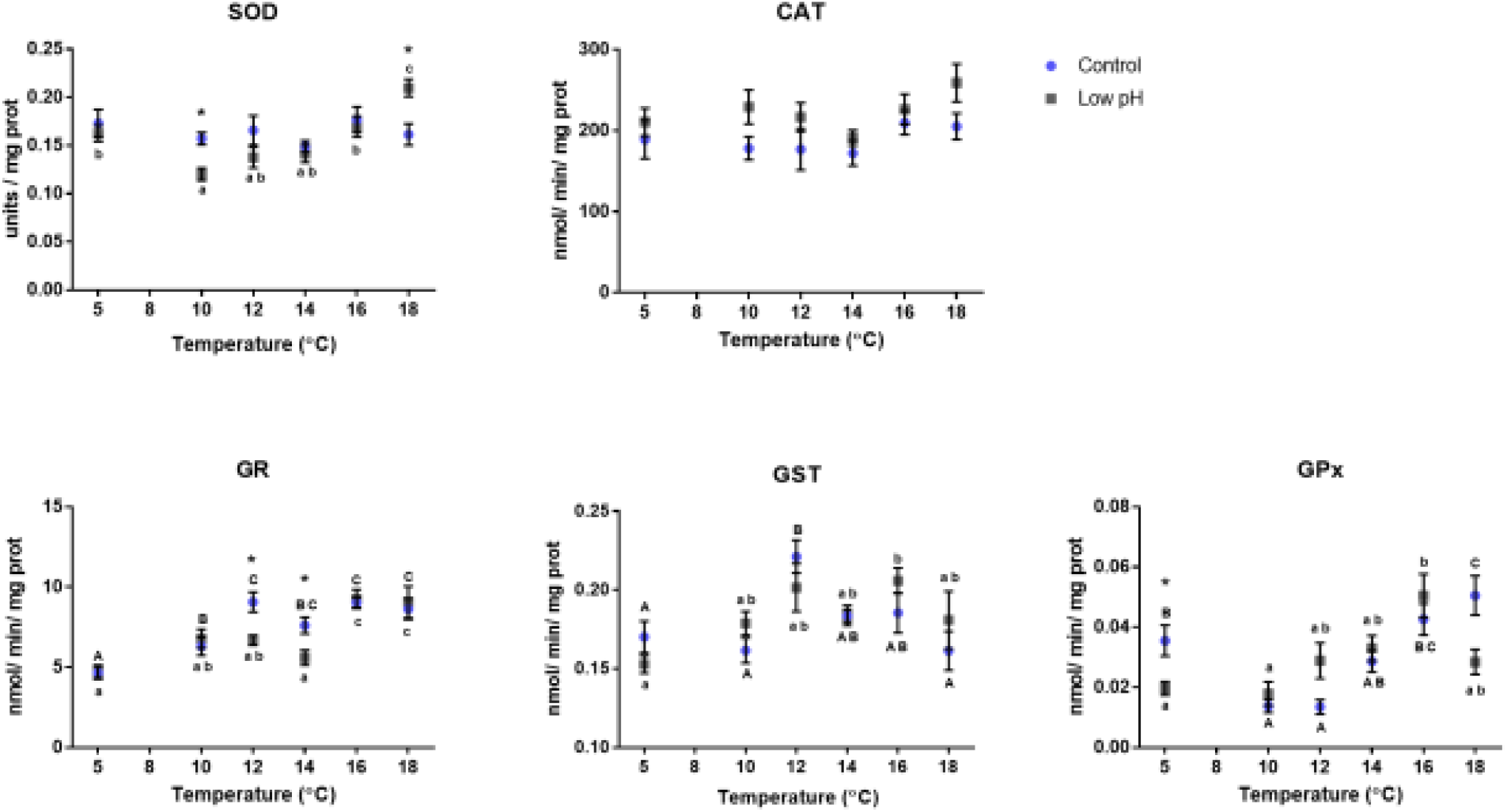
Oxidative stress indicators measured in liver of Atlantic halibut at different temperatures. From top left to right down: superoxide dismutase (SOD), catalase (CAT), glutathione reductase (GR), glutathione S-transferase (GST), glutathione peroxidase (GPx). Elevated CO_2_-treated groups significantly different from their respective controls are indicated by *. Letters indicate significant differences between temperatures within the same CO_2_ treatment. High and low caption letters indicate respectively control and elevated CO_2_ values. Error bars presented as ± SEM and confidence interval of 95 %.

**Figure 2.**
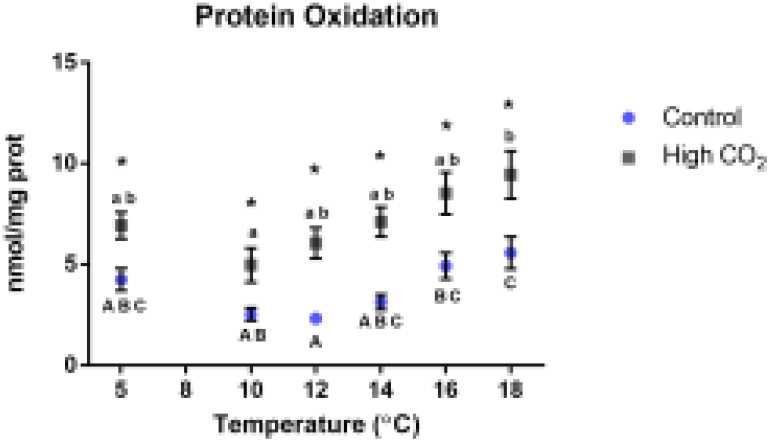
Oxidative damage measured as protein carbonyls, in liver of Atlantic halibut at different temperatures. Elevated CO_2_-treated groups significantly different from their respective controls are indicated by *. Letters indicate significant differences between temperatures within the same CO_2_ treatment. High and low caption letters indicate respectively control and elevated CO_2_ values. Error bars presented as ± SEM and confidence interval of 95 %.

**Figure 3.**
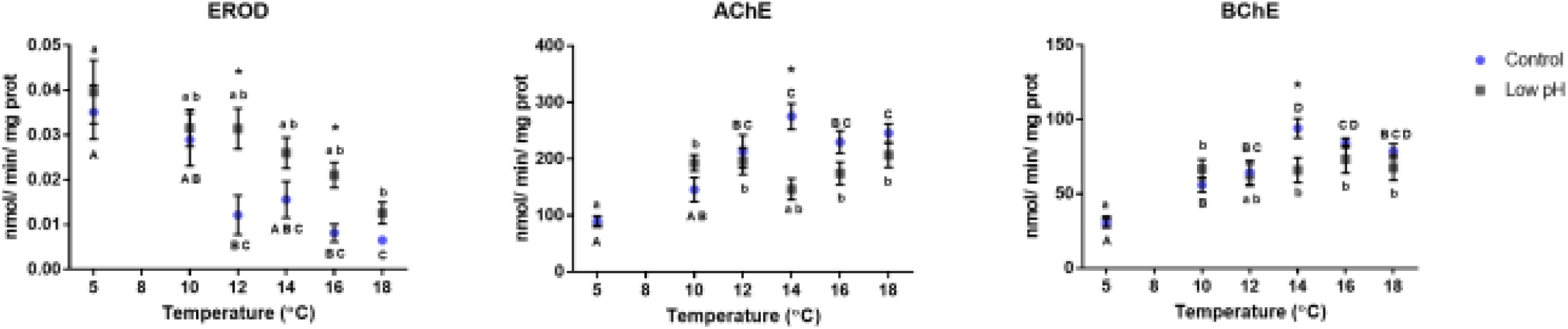
Biomarker responses, in liver of Atlantic halibut at different temperatures. Elevated CO_2_-treated groups significantly different from their respective controls are indicated by *. CYP1A activity measured as ethoxyresorufin O-deethylase (EROD), acetylchominesterase (AChE) and butyrylcholinesterase (BChE). Letters indicate significant differences between temperatures within the same CO_2_ treatment. High and low caption letters indicate respectively control and elevated CO_2_ values. Error bars presented as ± SEM and confidence interval of 95 %.

The elevated CO_2_ treatment resulted in reduced activity of SOD, except at 18 °C where there was a shift in this enzyme activity (higher activity in the elevated CO_2_ group). CAT activity was slightly higher in the elevated CO_2_-treated group. Both enzymes showed some changes related to temperature: SOD and CAT correlate significantly (*p* = 0.017) with one another in control groups, but not in elevated CO_2_-exposed fish. See Figure 1.

EROD activity and PC levels were consistently increased in the elevated CO_2_-treated groups at all temperatures, showing a clear CO_2_ effect. EROD activity (Fig. 3) was higher at lower temperatures and reduced with temperature increase in both control and elevated CO_2_-treated groups, while PC levels (Fig. 2) increased from temperatures of 10 to 18 °C in both control and elevated CO_2_-treated groups. The activities of GR, GST, and GPx varied within temperature treatments with a trend towards increasing with temperature (Fig. 1).

Both AChE and BChE increased significantly with increasing temperature, and while BChE was not affected by elevated CO_2_ levels, AChE was found to differ between these treatments. In addition, a significant interaction between temperature and CO_2_-treatment was identified in results from AChE measurements.

The two-way ANOVA revealed significant overall effects of temperature (Wilks’ Lambda = 0.048, F = 5.173, *p* < 0.001), pH (WL = 0.35, F = 10.96, *p* < 0.001) and an interaction between the two (WL = 0.321, F = 1.55, *p* < 0 0.021) measured parameters. Values for each specific parameter are listed in Table 1.

**Table 1.** Results from statistical testing of measurements conducted in tissue from halibut experimentally exposed to different temperatures and carbon dioxide concentrations. Values from the two-way ANOVA and Tukey’s post-hoc test investigating significant differences between the two treatment parameters as well as interactions between these two. Boldfaced text highlights significant differences. The total number of measurements (fish individuals) for each enzyme ranged from 72 to 77, and was 122 for protein carbonyls assay.

|  | Temperature |  | Elevated CO<sub>2</sub> |  | Temperature *<br>Elevated CO<sub>2</sub> |  |
| --- | --- | --- | --- | --- | --- | --- |
| Parameter | F | p | F | p | F | p |
| Superoxide dismutase (SOD) | 5.141 | **0.001** | 1.128 | 0.292 | 3.440 | **0.008** |
| Catalase (CAT) | 1.281 | 0.284 | 10.156 | **0.002** | 0.474 | 0.794 |
| Glutathione reductase (GR) | 12.423 | **0.000** | 1.697 | 0.198 | 1.783 | 0.130 |
| Glutathione S-transferase (GST) | 4.704 | **0.001** | 0.044 | 0.835 | 1.078 | 0.382 |
| Glutathione peroxidase (GPx) | 8.967 | **0.000** | 0.007 | 0.936 | 2.701 | **0.029** |
| Ethoxyresorufin O-deethylase (EROD) | 8.541 | **0.000** | 11.895 | **0.001** | 1.560 | 0.185 |
| Acetylcholinesterase (AChE) | 14.01 | **0.000** | 9.012 | **0.004** | 4.721 | **0.001** |
| Butyrylcholinesterase (BChE) | 16.07 | **0.000** | 3.324 | 0.072 | 2.127 | 0.071 |
| Protein carbonylation (PC) | 5.393 | **0.000** | 29.258 | **0.000** | 0.535 | 0.749 |

## Discussion

This study addresses the effects of elevated CO_2_ combined with different temperatures on biochemical responses in the Atlantic halibut. Our results are indicative of oxidative stress, which can result in damage macromolecules such as proteins (Pastore et al., 2003), as evident in increased protein carbonyl levels. Levels of oxidized proteins are often measured as protein carbonyls and these are frequently used as an indicator of oxidative stress (Carney Almroth et al., 2005; Dalle-Donne et al., 2003). Oxidation of proteins can lead to non-reversible conformational changes, which decrease enzyme activities and can result in protein degradation by proteases (Carney Almroth et al., 2005; Grune et al., 2004). We show that protein carbonyl levels in the elevated CO_2_ fish are consistently higher at all temperatures, and protein carbonyl levels are elevated at both high and low temperature extremes (Fig. 2). However, the antioxidant enzymes does not show the same pattern, suggesting that the exposure to elevated CO_2_ increased ROS formation, with consequent oxidative damage resulting from an insufficient antioxidant defense system.

The mechanisms underlying oxidative stress responses to ocean acidification (OA) are not fully understood, and three possible pathways have been described earlier in (Tomanek et al., 2011b), but induced oxidative stress and damage have been indicated as consequences of OA in several different groups of organisms (Hernroth et al., 2012; Kaniewska et al., 2012; Pimentel et al., 2015; Wood et al., 2016). We have previously shown that most of the Atlantic halibut used in our study were probably not experiencing acidosis or ionic homeostasis unbalance, since plasma lactate and ionic (K^+^, Na^+^, Ca^++^, Cl^-^) levels were the same in controls and elevated CO_2_ fish after 96 days of exposure (Bresolin de Souza et al., 2016). However, since we did not measure pH in the plasma or the intracellular compartment, we cannot determine whether the effects we measure are directly related to mitochondrial function or ROS production. Previous studies show that exposure to increased levels of CO_2_ can result in overall stress responses, changes in protein repair and degradation, as well as changes in the expression of hypoxia inducible factor 1 (HIF-1), a gene expressed in response to hypoxia stress (Dennis Iii et al., 2014). We hypothesize that the increased levels of protein carbonyls seen here may be related to the activation of hypoxia inducible factors (HIF-1) as well as the regulation of genes involved in oxygen transport and anaerobic energy production (Kassahn et al., 2009; Turrens, 2004). It has also been proposed that elevated CO_2_ exposure can affect the electron transport chain, increasing the production and release of ROS and other free radicals, (Dean et al., 1997; Tomanek, 2014a; Tomanek et al., 2011b).

The ideal growth temperature range for Icelandic Atlantic halibut juveniles (used in this study) is from 11 to 14 °C (Björnsson and Tryggvadóttir, 1996), while 18 °C is their upper tolerance limit and results in a reduced growth rate (Imsland, 2001; Langston et al., 2002). In overall, antioxidant enzyme activities did not increase linearly with increasing temperature, in agreement with previous studies in fish (Madeira et al., 2013; Vinagre et al., 2012). Oxidative stress-related enzymes increase in activity with increased temperature until reaching a peak, and after the temperature peak the enzymes activity decreases. The temperature of this peak is related to the thermal niche of the species and is species-specific (Gräns et al., 2014a).

Gräns et al. (2014a) studied effects of global climate change on Atlantic halibut: acclimation to warmer temperatures resulted in increased aerobic scope and cardiac performance, an effect that was even more pronounced by elevated CO_2_ exposure. These effects were not reflected in growth rate, which was slower at the warmest temperatures (16 and 18 °C), leading the authors to conclude that oxygen uptake was not a limiting factor for growth. However, this study also found no differences in oxygen consumption between fish kept at control or elevated CO_2_ (pH 8.1 or 7.7 respectively), at any given temperature. Therefore, we can conclude that increased oxygen consumption, potentially leading to an increase in ROS production, is not directly responsible for the increase in levels of protein carbonyls seen in the elevated CO_2_-treated fish in the current study.

Our results also showed that activity of GR was lower at 12 and 14 °C in the elevated CO_2_-treated groups, which could possibly indicate an overconsumption of glutathione in other cellular reactions. Glutathione is an important molecular antioxidant, and the regeneration of its reduced form from oxidized molecules is catalyzed by GR. GPx, which is responsible for reducing both hydrogen and lipid peroxides, protecting the cells against the damaging effects of lipid peroxidation (Winston and Di Giulio, 1991), was in general more affected by temperature, especially at the extremes (5 and 18 °C), than by elevated CO_2_. The activities of two additional antioxidant enzymes, SOD and CAT, correlate significantly with one another in control fish but not in elevated CO_2_-treated fish. A mismatch in SOD and CAT activities can result in an ineffective metabolism of ROS, allowing ROS to interact with other molecules, causing oxidative damage (Halliwell and Gutteridge, 1999).

CYP1A activity (EROD) was higher in the elevated CO_2_-treated fish in all studied temperatures, thus indicating that there is a CO_2_ effect on EROD activity, which is independent of xenobiotic exposure. The effects of pH on EROD activity are not clear, yet most studies to date have exposed fish to pH levels much lower than in the current study ((Whyte et al., 2000) and references therein). Research on the mechanisms of toxicity behind CYP1A induction show that EROD activity not only indicates chemical exposure but may also reflect effects of other abiotic factors (Whyte et al., 2000). Induction of CYP1A in fish requires the activation of cytosolic aryl hydrocarbon receptors (AhR), followed by the transcription of the Ah-gene battery and subsequent synthesis of proteins, including CYP1A and many phase II enzymes (Whyte et al., 2000). Hence, the observed CO_2_-dependent increase of EROD activity in the current study cannot be due to a xenobiotic induction, but rather a post-translational or kinetic regulation of the enzyme. In addition, EROD had a strong negative correlation with temperature, in contrast to antioxidant enzyme activities measured here, indicating that these proteins may have different stability or regulatory pathways linked to temperature (Regoli and Giuliani, 2013; Solé et al., 2015). Temperature impacts the composition and hence the fluidity of membranes ((Nikinmaa, 2013) and references therein), so this may have a greater impact on EROD, a membrane bound protein, than on the cytosolic antioxidant enzymes. In addition, it is possible that cell membrane properties are influenced by oxidative stress-dependent lipid peroxidation, changing membrane fluidity and the integrity of biomolecules associated with membranes (Carney Almroth et al., 2005).

In this study, we also measured effects in activity of two cholinesterases, AChE which is known for its role in neurotransmission via hydrolysis of acetylcholine, and BChE, suggested to play a role in ghrelin regulation and aggression (Brimijoin et al., 2016; Chen et al., 2015). AChE occurs mostly in brain, neurons and muscle but is also present in many other tissues, while BChE predominates in liver, lungs, plasma, and neuroglia (Chen et al., 2015; Pohanka, 2014). Both enzymes have been used as indicators for xenobiotic exposure (Sanchez et al., 2011; Sturm et al., 1999). AChE has been shown to be sensitive to stress and inflammation (Bresolin de Souza et al., 2014b; Ming et al., 2015). In addition, the activity of serum BChE is an indicator of systemic inflammation in humans, and the more severe the inflammation, the smaller the concentration of serum BChE (Zivkovic et al., 2015). In the present study, both AChE and BChE show a positive correlation with temperature, but this tendency was less evident in the elevated [CO_2_]-treated fish. However, current understanding of the effects of temperature on AChE and BChE activities are conflicting. Some studies show positive effects and others negative effects, while still others find no significant differences ((Solé et al., 2015) and references therein). Interestingly, both AChE and BChE were decreased at elevated [CO_2_] treatment at 14 °C, which is within the optimal growth temperature range for this species (Gräns et al., 2014b) but outside normal water pH levels. We would also like to propose that further investigation into BChE and its role in physiological responses to environmental stressors could prove interesting since: BChE has been indicated as playing a role in ghrelin regulation and aggression in knockdown mice (Chen et al., 2015); ocean acidification can result in decreased BChE activity as shown here; and previous studies have shown behavioural changes in stickleback when exposed to increased [CO_2_], including effects on boldness and exploratory behaviour (Jutfelt et al., 2013; Näslund et al., 2015).

Metabolic acclimations to deal with changes in environmental temperature have been previously described (Pörtner et al., 2005; Somero, 2012). Arrhenius’ law dictates that acclimation to lower temperatures includes increased protein synthesis to compensate for lower reaction rates (Solé et al., 2015; Whyte et al., 2000). This mechanism could explain the negative relationship of EROD activity and temperature in our study, which may be due to increased amounts of CYP1A at lower temperatures, and vice-versa. There are previous studies with fish showing the same negative correlation between EROD activity and temperature acclimation (Solé et al., 2015; Whyte et al., 2000).

Ocean acidification is thought to raise metabolic rates in aquatic organisms in order to supply the cells with additional energy to cope with the physiological changes caused by such environmental variations (Nikinmaa, 2013). In addition, when animals are close to their thermal limits, highest or lowest preferred temperature tolerated, even a small disturbance (small rise in temperature) may reduce their activity scope and consequently reduce ecological success (Nikinmaa, 2013). The extra energetic cost associated with these processes is expected to result in an increase oxidative stress (Tomanek, 2014a). Evidences of metabolic changes resulting from exposure to elevated CO_2_ are found in these experimental fish in a previous study (Bresolin de Souza et al., 2014b), seen in the modulation of proteins such as glyceraldehyde 3-phosphate dehydrogenase, fructose-1,6-phosphate aldolase, and malate dehydrogenase. Similar changes regarding higher expression of metabolic enzymes and increased oxidative stress have been shown in the Pacific oyster (*Crassostrea gigas*) exposed to elevated CO_2_ (Timmins-Schiffman et al., 2014). Oxidative stress is a co-stress of temperature and elevated CO_2_ low pH, and stress-mediated ROS can lead to shifts in energy metabolism. This can be accompanied by the activation of pathways of ATP production; excess ROS and shifts in energy metabolism might impact protein homeostasis through e.g. protein denaturation or lack energy for protein synthesis and normal function (Tomanek, 2011b; Tomanek, 2014b).

## Conclusion

This study provides insights regarding how fish are affected by the combined stress of elevated CO_2_ and temperature. We show physiological changes, possibly related to detrimental effects and/or acclimation mechanisms, that indicate impacts of future climate change, as modelled in the conditions in our experiments. Here we show that elevated CO_2_ exposure can induce oxidative stress, evident in accumulation of protein carbonyls. In addition activities of esterases and detoxification enzymes were shown to be affected by temperature and CO_2_ levels. The results presented here support the hypothesis that CO_2_ levels estimated to occur at the end of this century could pose physiological challenges to marine fish.

## Acknowledgements

We thank the GRIP project members for their collaboration and support, and Niklas Dahr for technical assistance. The study was supported by the University of Gothenburg Platform for Integrative Physiology (GRIP) and the Swedish Research Council for Environment, Agricultural Sciences and Spatial Planning (FORMAS).

